# Experience-dependent neuroplasticity of the developing hypothalamus: integrative epigenomic approaches

**DOI:** 10.1101/191791

**Authors:** Annie Vogel Ciernia, Benjamin I. Laufer, Keith W. Dunaway, Charles E. Mordaunt, Rochelle L. Coulson, Theresa S. Totah, Danielle S. Stolzenberg, Jaime Frahm, Akanksha Singh-Taylor, Tallie Z. Baram, Janine M. LaSalle, Dag H. Yasui

**Affiliations:** Department of Medical Microbiology and Immunology, University of California, Davis, CA, USA; Department of Psychology, University of California, Davis, CA, USA; Center for Comparative Medicine, University of California, Davis, CA, USA; Present Address: Active Motif, 1914 Palomar Oaks Way # 150, Carlsbad, CA, USA; Department of Pediatrics and Anatomy/Neurobiology, University of California, Irvine, CA, USA; UC Davis Genome Center, UC Davis, Davis, CA, USA; UC Davis MIND Institute, UC Davis, Davis, CA, USA

**Keywords:** Augmented maternal care, epigenomics, DNA methylation, early-life stress, hypothalamus

## Abstract

**Background:** Maternal care during early-life plays a crucial role in the sculpting of the mammalian brain. Augmented maternal care during the first postnatal week promotes life-long stress resilience and improved memory compared with the outcome of routine rearing conditions. Recent evidence suggests that this potent phenotypic change commences with altered synaptic connectivity of stress sensitive hypothalamic neurons. However, the epigenomic basis of the long-lived consequences is not well understood.

**Methods:** Here, we employed whole-genome bisulfite sequencing (WGBS), RNA-sequencing (RNA-seq), and a multiplex microRNA (miRNA) assay to examine the effects of augmented maternal care on DNA cytosine methylation, gene expression, and miRNA expression.

**Results:** A significant decrease in global DNA methylation was observed in offspring hypothalamus following a week of augmented maternal care, corresponding to differential methylation and expression of thousands of genes. Differentially methylated and expressed genes were enriched for functions in neurotransmission, neurodevelopment, protein synthesis, and oxidative phosphorylation, as well as known stress response genes. Twenty prioritized genes with three lines of evidence (methylation, expression, and altered miRNA target) were identified as highly relevant to the stress resiliency phenotype.

**Conclusions:** This combined unbiased approach enabled the discovery of novel genes and gene pathways that advance our understanding of the central epigenomic mechanisms underlying the profound effects of maternal care on the developing brain.

## Background

Early postnatal care in mammals is critical for survival as well as neurodevelopment. Specifically, early maternal care can epigenetically alter the behavior of offspring for their entire lifetime. Rodent models manipulating maternal care and maternal stress have been used to examine the molecular and cellular mechanisms underlying the long-term behavioral changes observed in the offspring^1–3^. In the augmented maternal care (AMC) paradigm, daily separation of rat dams from their pups for 15 minutes from postnatal day 2 (PND2) to postnatal day 8 (PND8) triggers the increased maternal behaviors of arched-back nursing, as well as elevated licking and grooming of the pups by the dam^3,4^. Subsequently, it was found that multiple days of pup handling within the first postnatal week of life were required for the AMC response, suggesting AMC initiates changes during this critical period of neurodevelopment in the offspring^5,6^. The repeated, daily sessions of enhanced maternal care triggered in the AMC paradigm result in offspring that show reduced plasma corticosterone levels in response to an acute restraint stress^7–11^, reduced anxiety related behaviors^11,12^, resilience to depressive-like behaviors^11^, and enhanced performance on several cognitive tasks as adults^12,13^.

AMC appears to alter brain function for the lifetime of the animal^14^ and as adults AMC offspring show hypo-activation of the HPA axis in response to stress. Previous work has demonstrated that this mechanism depends upon temporally coordinated regulation of both corticotrophin releasing hormone (CRH) in the paraventricular nucleus (PVN) of the hypothalamus^5,13^ and glucocorticoid receptor (GR, encoded by *Nr3c1*)^6^ in the hippocampus^13,15^. *Crh* transcripts in the hypothalamus are significantly reduced immediately following AMC at PND9^5,6^ and remain repressed throughout adulthood^9^. In comparison, increased *Nr3c1* expression in the hippocampus was delayed until PND23^9^, suggesting that early gene expression changes in the PVN of the hypothalamus drive the initial molecular mechanisms underlying AMC. Together, the elevated hippocampal *Nr3c1* expression and reduced hypothalamic *Crh* expression^2,5,11,16^ in AMC adult offspring suggest that increased negative feedback signaling may underlie the observed hypo-responsive HPA axis in the adult AMC offspring.

These studies have raised an important question about how hypothalamic reprogramming is initiated and maintained by AMC induced early-life gene expression changes. AMC initiates transient structural changes in CRH-expressing cells in the hypothalamus of pups^12,16^. By PND9 there is decreased excitatory synaptic drive onto hypothalamic *Crh* expressing neurons in AMC offspring, that returns to baseline by adulthood (PND30-45)^16^, suggesting that the AMC induced reduction in *Crh* expression is maintained long-term by additional mechanisms^14^. *In vitro* studies have suggested that the synaptic changes initiate transcriptional regulation in these CRH positive cells^12^. Specifically, recent work has implicated the neuron restrictive silencing factor (NRSF) in establishing long-term epigenetic gene expression patterns resulting from AMC^14,17^.

NRSF expression and binding to the *Crh* gene is increased in hypothalamus of AMC offspring at PND9 but is no longer present in adulthood^12,16^. Thus, there is little information about the comprehensive epigenomic processes by which augmented sensory signals from the mother result in enduring, large-scale reprogramming of gene expression in the hypothalamus. In hippocampus, prior studies have described epigenetic DNA methylation alterations and H3K9 acetylation in response to differences in maternal care at *Nr3c1*^18^. The search for additional genes underlying AMC effects on adult hippocampus revealed alterations to gene expression of more than 900 genes related to cellular metabolism and energy production, signal transduction, protein synthesis (mainly ribosomal genes), and neurodevelopment^19^. Targeted analysis of a seven megabase (Mb) region of chromosome 18 flanking *Nr3c1* revealed that the clustered protocadherins (*Pcdh*) showed the highest differential response in DNA methylation^20^.

Together, these combined results indicate that NRSF binding in PND9 hypothalamus ^12,16^ as well as differential gene expression and DNA methylation in adult hippocampus are involved in regulating long-term behavioral changes induced by AMC. However, whereas NRSF ChIP-seq has identified gene sets contributing to AMC-induced cellular changes^12^, a multi-methodology genome-wide approach has not been undertaken in the hypothalamus^21^. To address this gap in knowledge we performed an integrated, genome-wide examination of epigenetic and gene expression differences in hypothalamus from PND9 pups in response to AMC. These experiments identified that the response to AMC occurs not only in the genes previously identified such as *Crh*, but in thousands of genes throughout the genome. In addition to a significant global reduction in hypothalamic DNA methylation levels, AMC induces differential methylation and expression in mRNA transcripts related to neurodevelopment, synaptic signaling, ribosome function, and cellular stress. The results provide strong evidence for the role of epigenomic modifications in regulating gene expression profiles related to resilience and offer a suite of prioritized genes and gene pathways for the development of diagnostic biomarkers and therapies for neuropsychiatric disorders in humans.

## Methods

Detailed methods are provided in the **Supplemental Information** section. All animal studies were performed in accordance with UC Davis IACUC protocol #18745. Augmented Maternal Care (AMC) was performed as described previously^13,16^. Briefly, pups were gathered at birth and five male and five female pups were randomly assigned to dams. Pups were removed for 15 min at 9 AM each day from PND2-PND9 to a separate cage in a different room equipped with a heating pad. Further details can be found in the **Supplemental Information (SI) Materials and Methods**. For WGBS DNA methylation analysis, genomic DNA was converted by bisulfite treatment using a Zymo kit followed by Illumina sequencing library preparation. WGBS libraries were sequenced on an Illumina Hi-Seq 2500. DMR identification and PMD analysis was performed as detailed in Schroeder^22^ and Dunaway^23^. The **SI Materials and Methods** section contain an in depth description of WGBS methods. Bisulfite converted genomic DNA was amplified using custom, gene specific primers and amplicons were analyzed by pyrosequencing on a Biotage PSQ-96MA Pyrosequencer. A detailed description of pyrosequencing methods is contained in the **SI Materials and Methods**. For RNA-seq total RNA was prepared from tissue using a Qiagen RNeasy Kit (Qiagen). Total RNA was depleted of ribosomal RNA for creation of stranded RNA-seq libraries (**See SI Materials and Methods**). Bar coded RNA-seq libraries were sequenced on an Illumina Hi-Seq 2500 platform. Analysis of RNA-seq reads is described in **SI Materials and Methods**. Validation for differential RNA-seq identified genes was performed on samples from the same cohort of PND9 male and female hypothalami (**SI Materials and Methods**). For miRNA transcript analysis a multiplex miRNA assay of miRNA transcripts was performed using an nCounter Rat miRNA expression assay. A detailed description of miRNA analysis is available in **SI Materials and Methods**. All sequencing data will be available on the NIH GEO database.

## Results

### AMC induces large-scale hypo methylation in the PND9 hypothalamus

To investigate genome-wide DNA methylation changes associated with AMC, hypothalami from AMC and control male rat pups were chosen for whole-genome bisulfite sequencing (WGBS) based on significant (*p*=0.0042) differential AMC (total licking and grooming) at postnatal day 9 (**Supplemental Figure 1**). Principal component analysis (PCA) was performed on 20,000 base (20-Kb) windows of percent DNA methylation with CpG islands masked (Figure 1A) to explore genome-wide patterns of methylation between treatment conditions in the individual rats. Methylation levels in individual PND9 hypothalami were separated by AMC condition for principal component 1 (30.2% of variance explained), indicating differences in methylation levels between conditions. Unexpectedly, hypothalami from AMC pups displayed significant global hypomethylation (*p*=0.015) at PND 9 (Figure 1B). Specifically, global CpG methylation levels for augmented hypothalami averaged 76.08% while controls had levels of 79.20% yielding to a 3.12% reduction **(Supplemental Table 1**). Non-CpG methylation levels in augmented and control hypothalami ranged from 0.86 to 1.46% of total cytosines (**Supplemental Table 1**) consistent with previous findings of normal neurodevelopment^24^. CpG hypomethylation was observed across all chromosomes with the exception of chromosome 13 and X, indicating a genome-wide impact of AMC on CpG methylation levels (Figure 1C). Interestingly, windowing over different genomic features revealed hypomethylation specifically in the region 5 kb upstream of the transcriptional start site (TSS) of CpG island promoters (*p*=0.0057) in AMC hypothalamus (**Supplemental Figure 2**). To identify potential hypothalamic genes and pathways contributing to AMC, large-scale differential DNA methylation patterns were identified from the WGBS data sets. Partially methylated domains (PMDs) were mapped from AMC and control WGBS data sets using a hidden Markov model (HMM) previously described for detecting partially methylated domains in neuronal cell lines^22^. AMC PMDs averaged 7.3% of the genome and control PMDs averaged 6.7% of the genome.

**Figure 1.**
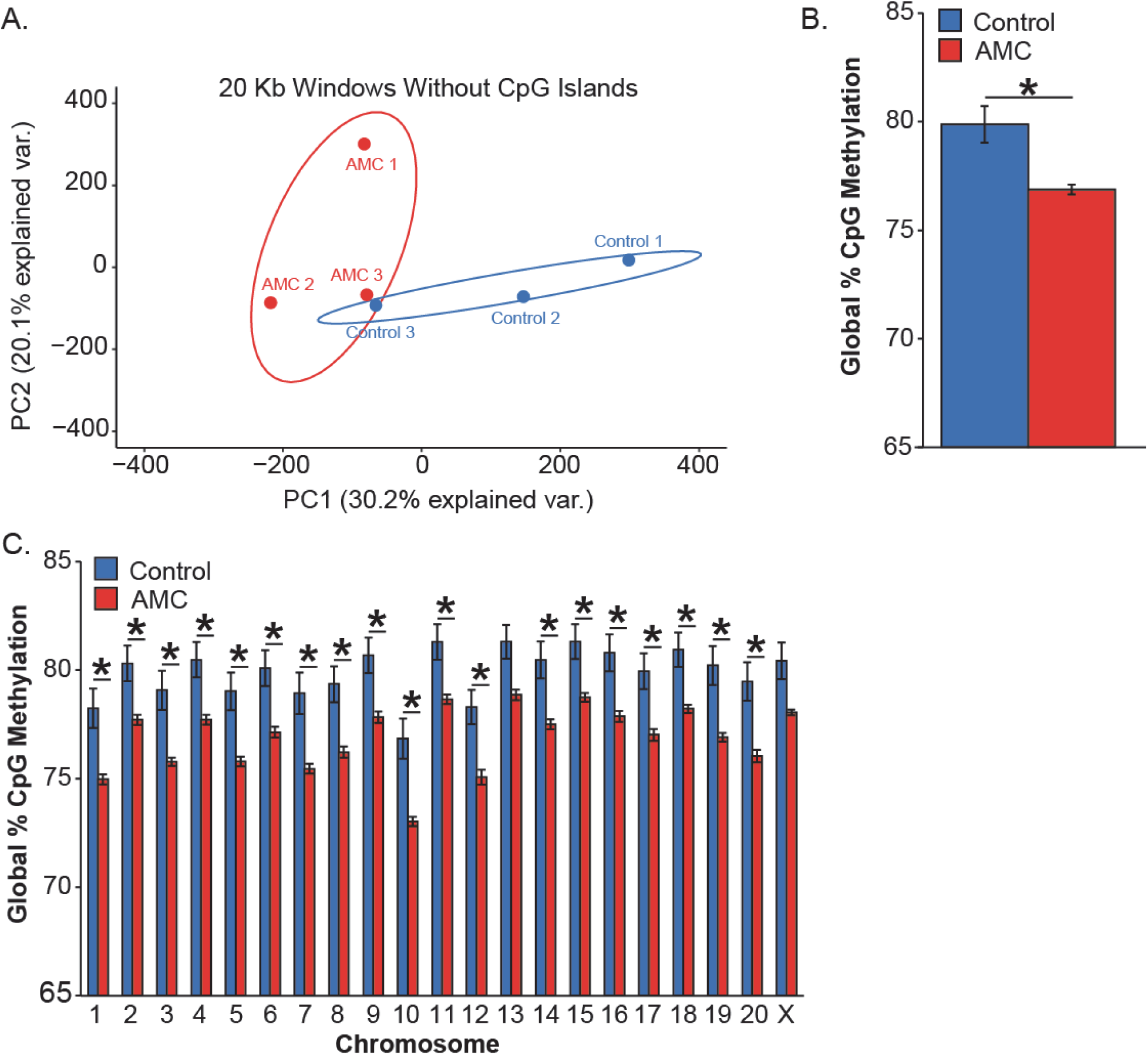
Reduced DNA methylation in hypothalamus is associated with AMC. (A) Principal components 1 and 2 of the average DNA methylation in 20 kb windows with masking of CpG Islands separately groups AMC samples (red outline, *n=3*) from control samples (blue outline, *n=3*). Outlines represent 95% confidence intervals. (B) Average genome wide CpG methylation is significantly reduced (*p*=0.015) by 3.12% in AMC samples (*n=3*) compared with control samples (*n=3*). (C) DNA hypomethylation in the AMC hypothalami at PND9 was significant (\**p*<0.05) on every chromosome except chromosome 13 and X (*p*<0.52 and 0.054 respectively). RM ANOVA AMC group x chromosome interaction F (20.80) = 21.89, *p*<0.0001; * Benjamini-Hochberg corrected t-tests for each chromosome all *p*<0.05.

Although consistent with genome-wide hypomethylation the increased PMD coverage with AMC was not significant (*p*=0.158) (**Supplemental Table 2**). While PMDs were found throughout the genomes of both groups, only five PMDs were differential by treatment group (Table 1). Consistent with genome-wide hypomethylation shown in Figure 1B, these five PMDs were found in AMC but not control hypothalami. AMC PMDs covered 674,924 bases and overlapped with seven genes. These PMD genes included *Dbn1*, *Prr7*, *Foxo3*, and *Cnksr2,* which function in neuronal development, and *Pak6* which functions in the MAP kinase pathway to regulate cytoskeletal dynamics underlying multiple brain processes^25–29^(Table 1). AMC PMDs also contained *AY383691* and *Ankrd63*, which are gene products without well-established functions.

**Table 1.** Gene transcripts located within AMC PMDs.

| size (bases) | genomic position | genes | PMD | function |
| --- | --- | --- | --- | --- |
| 24,868 | chr17:9,703,631-9,678,763 | *Dbn1* | AMC | Cell migration, extension of neuronal processes and plasticity of dendrites |
|  |  | *Pr77* | AMC | Enriched in post-synaptic densities and dendritic spines |
| 4,838 | chr20:46,425,344-46,430,182 | *Foxo3* | AMC | Transcriptional activator which triggers apoptosis in the absence of survival factors and during oxidative stress |
| 11,664 | chr3:110,484,414-110,496,078 | *Ankrd63* | AMC | Ankyrin repeat domain protein 63 |
|  |  | *Pak6* | AMC | cytoskeleton rearrangement, apoptosis, and the mitogen-activated protein (MAP) kinase signaling pathway |
| 168,184 | chrX:39,545,119-39,713,303 | *Cnksr2* | AMC | assembly of synaptic proteins at the membrane and coupling of signal transduction to membrane/cytoskeletal remodeling |
| 471,627 | chr5:35,190,215-35,661,842 | *AY383691* | AMC | unknown |

### AMC DMRs are linked to neurodevelopmental signaling and neurotransmission

While PMDs highlight broad regions of differential methylation associated with AMC, few genes were found within these loci. Therefore to identify small scale (<3 Kb) methylation changes, differentially methylated regions (DMRs) were identified from WGBS data^23,30^. With this bioinformatic approach, a total of 9,439 DMRs between AMC and Control PND9 hypothalami were identified (**Supplemental Table 3**). Of these 9,439 AMC DMRs, 6,228 had reduced methylation levels in the AMC samples, which is consistent with the genome-wide hypomethylation shown in Figure 1. To investigate the relationship between large-scale PMDs and small-scale DMRs, the two genomic features were compared. Surprisingly, of the 9,439 DMRs identified, only one in *Pak6* was found within the five AMC PMDs. Visualization of the DMRs using hierarchical clustering analysis (HCA) of significant (*p*<0.05) DMRs from hypothalami clustered each rat by treatment (Figure 2A). Both PCA of 20kb windows (Figure 1A) and HCA (Figure 2A) independently partitioned the AMC and control hypothalami into separate groups, indicating that distinct methylation patterns distinguish the AMC from control hypothalamic samples at two different resolutions. Motif enrichment analysis of the DMRs revealed a significant (*q*<0.0001) enrichment only for CTCF binding sites, which were present in 4% of DMRs (Figure 2B). CTCF binds these sites in a methylation sensitive manner and functions as a genomic insulator critical to chromatin architecture^31^. Of the 9,439 total DMRs identified, 5,284 were between 5 kb upstream and 1 kb to downstream of 4,023 coding and non-coding RefSeq genes in the rat genome (Rn6) (**Supplemental Table 3**). Forty-four percent of the DMRs were intergenic and 35.3% were localized to introns. Fourteen percent of DMRs were immediately upstream of genes and only 0.4% and 1.1% were located in the 5’ and 3’ prime untranslated regions (UTRs) respectively (**Supplemental Figure 3A**). For the DMRs annotated to within genes, the majority (93.6%) were localized to coding genes and 4.1% were located in long non-coding RNAs (lncRNAs) (**Supplemental Figure 3B**).

**Figure 2.**
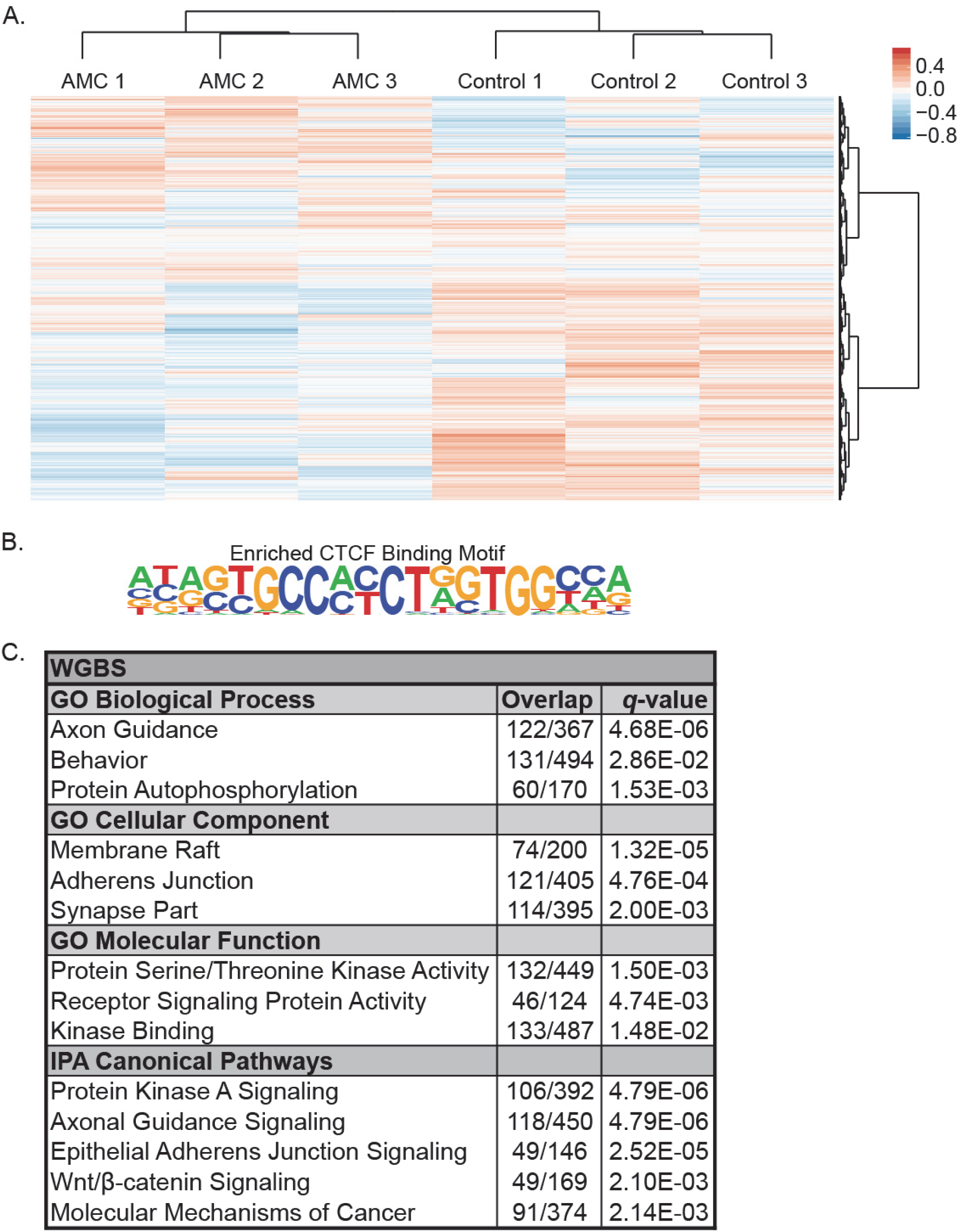
Differences in DNA methylation between AMC (*n*=3) and control (*n*=3) P9 hypothalami. (A) Hierarchical clustering of significant (*p*<0.05) DMRs. Individual percent methylation values are normalized to the mean of each DMR. (B) CTCF motif binding motif was significantly (*q*<0.0001) enriched in DMRs with 4.24% of DMRs containing the binding motif compared to 3.15% of background sequences with the motif (*q*<0.00001). (C) Significant (*q*<0.05) enriched gene ontologies and pathways of genes mapped to significant (*p*<0.05) DMRs. Overlap corresponds to genes observed compared with the total genes in the pathway.

To comprehensively identify the functions associated with the 4,023 genes mapped to AMC DMRs, gene ontology (GO) and pathways analyses were performed (Figure 2C). Gene ontology analysis revealed a significant (*q*<0.05) enrichment for the biological processes of axon guidance, behavior, and protein auto-phosphorylation. Membrane raft, adherens junction, and synapse part were the significantly (*q*<0.05) enriched cellular components. Protein serine/threonine kinase activity, receptor signaling protein activity, and kinase binding were the most unique significant (*q*<0.05) molecular functions. Pathway analyses revealed a significant (*q*<0.05) enrichment for protein kinase A signaling, axonal guidance signaling, epithelial adherens junction signaling, Wnt/β-catenin signaling, and molecular mechanisms of cancer. Notably, the molecular mechanisms of cancer pathway are a collection of developmental signaling pathways. Overall, all of the above GO terms and pathways are related to neurodevelopment as well as synaptic and cellular signaling (Figure 2C).

To further examine the DMRs for relevance to AMC, the 9,439 DMRs were subject to permutation testing, which yielded seven high confidence or “gold” DMRs (family-wise error rate (FWER) < 0.05) (**Supplemental Table 4**). These gold DMRs were annotated to *LOC00911370*, *Tuba3b*, *Glb1l*, *Papolb*, *Grb10*, *Tenm3,* and *Zfp332a* (**Supplemental Table 4**). *Zfp322a* and *Tenm3* both exhibited hypomethylation and increased gene expression in AMC pups. Interestingly, *Tenm3* plays a critical role in axon guidance and *Zfp322a* is a transcription factor that inhibits cellular differentiation^32,33^. The hypomethylation of the *Tenm3* associated DMR was validated (*p*=0.01) by an independent pyrosequencing assay (**Supplemental Figure 4**).

### Differential AMC gene expression influences translation and oxidative phosphorylation

To investigate the regulatory relationship between DMRs and gene expression, RNA-seq analysis was performed on postnatal day 9 (PND9) male hypothalami (Figures 3A and B & **Supplemental Table 5**). Examination of the top 500 most variably expressed genes revealed clustering by AMC condition (Figure 3A). For differential analysis, a total of 2,464 significant (*q*<0.05) differentially expressed (DE) transcripts were identified between AMC and control hypothalami (Figure 3B, 3C & **Supplemental Table 6A & 6B**). There were 1,411 genes with reduced expression and 1,053 with increased expression following AMC. Log_2_ fold changes in gene transcripts ranged from 2.67 to - 2.08 (**Supplemental Table 6A & 6B**) and the top 20 significant (*q*<0.05) genes ranked by fold change are shown in Table 2. Consistent with previous studies that have established a key role for stress-induced neuropeptide *Crh*^5,11^, we also found that *Gal*, *Oxt,* and *Ucn3,* transcripts encoding galanin, oxytocin and the Crh homologue urocortin, respectively, were significantly reduced in handled pup hypothalamus as were transcripts for the *Crh1* receptor (*p*<0.05)(Table 2 & Figure 3B).

**Figure 3.**
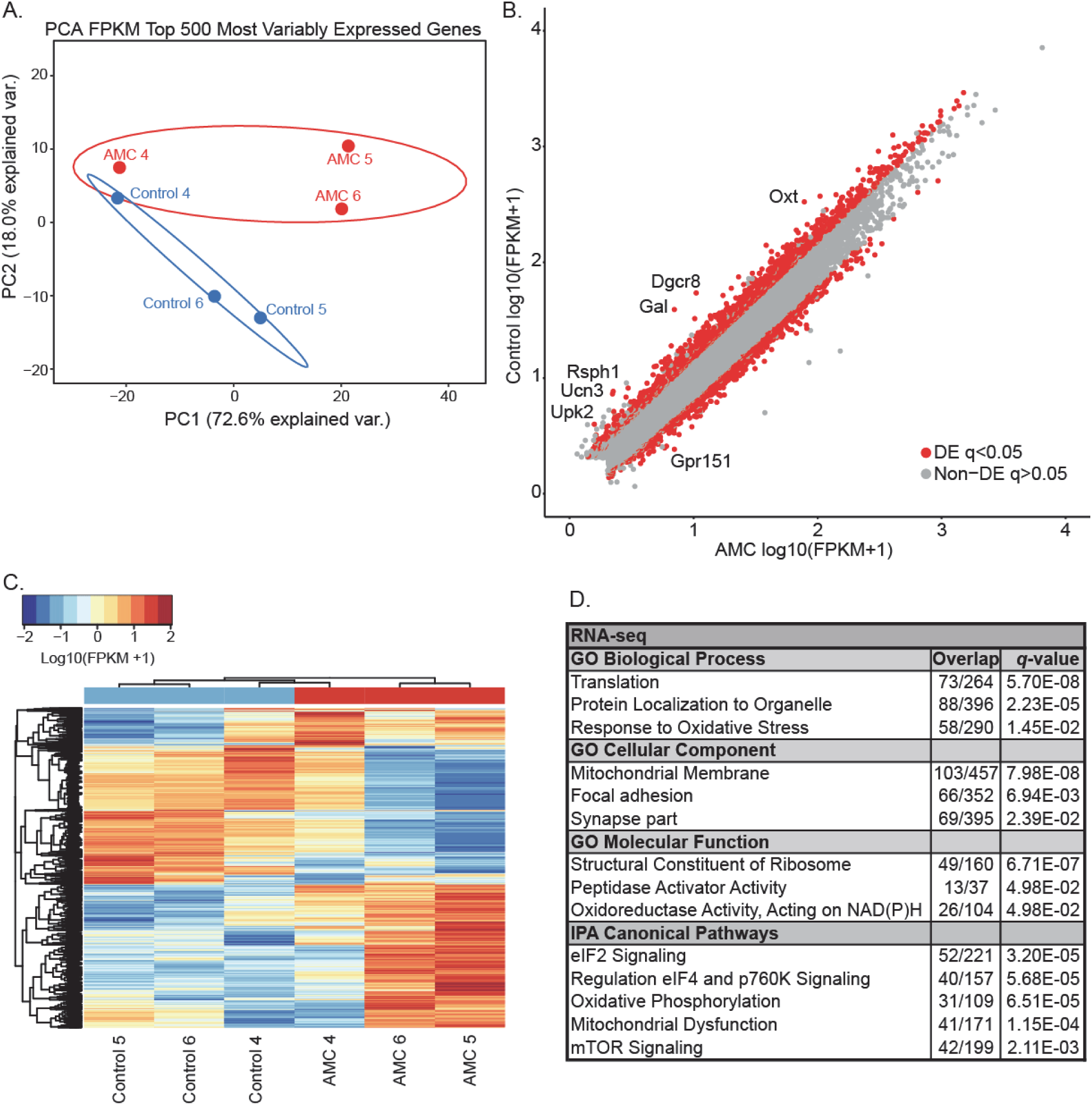
Differences in gene expression between AMC (*n*=3) and control (*n*=3) PND9 hypothalami. (A) Principal components analysis (PCA) of the top 500 most variably expressed genes. Outlines represent 95% confidence intervals. (B) Scatter plot DE transcripts with the top DE transcripts labelled. Grey indicates no significant difference in expression, while red indicates a significant (*q*<0.05) difference. (C) Hierarchal clustering analysis (HCA) of significant (*q*<0.05) DE transcripts. (D) Significant (*q*<0.05) enriched gene ontologies and pathways from significantly (q<0.05) DE transcripts. (E) Most significant (*q*<0.0001) transcription factor binding site identified by motif enrichment analysis.

**Table 2.** The top 20 elevated and repressed AMC RNA transcripts.

| Gene Symbol | q-value | log2 FC | Gene Name(s) |
| --- | --- | --- | --- |
| *Gpr151* | 0.002 | 2.1 | G Protein-Coupled Receptor 151 / Galanin Receptor 4 |
| *Ebf1* | 0.002 | 1.7 | Early B-cell Factor 1 |
| *Rbm46* | 0.002 | 1.7 | RNA Binding Motif Protein 46 |
| *Stk17b* | 0.002 | 1.7 | Serine/Threonine Kinase 17b |
| *Snhg4* | 0.006 | 1.6 | Small Nucleolar RNA Host Gene 4 |
| *Dcun1d1* | 0.002 | 1.6 | Defective In Cullin Neddylation 1 Domain Containing 1 |
| *Zfp347* | 0.002 | 1.6 | Zinc Finger Protein 347 |
| *Slc5a7* | 0.002 | 1.6 | Solute Carrier Family 5 Member 7 / Choline Transporter |
| *Rgs4* | 0.003 | 1.5 | Regulator Of G-Protein Signaling 4 / Schizophrenia Disorder 9 |
| *Pabpc2* | 0.008 | 1.5 | Poly(A) Binding Protein, Cytoplasmic 2 |
| *Sh3bgrl* | 0.002 | 1.5 | SH3 Domain Binding Glutamate Rich Protein Like |
| *Lpar4* | 0.002 | 1.5 | Lysophosphatidic Acid Receptor 4 |
| *Tlr4* | 0.005 | 1.5 | Toll Like Receptor 4 |
| *Rybp* | 0.002 | 1.5 | RING1 And YY1 Binding Protein |
| *Thoc2* | 0.002 | 1.5 | THO Complex 2 / Mental Retardation, X-Linked 12/35 |
| *Mier1* | 0.002 | 1.5 | Mesoderm Induction Early Response 1, Transcriptional Regulator |
| *Wfdc1* | 0.009 | 1.5 | WAP Four-Disulfide Core Domain 1 / Ps20 Growth Inhibitor |
| *Stag2* | 0.002 | 1.5 | Stromal Antigen 2 / Cohesin Subunit SA-2 |
| *Arrdc3* | 0.002 | 1.4 | Arrestin Domain Containing 3 |
| *Slc6a4* | 0.002 | 1.4 | Solute Carrier Family 6 Member 4 / Serotonin Transporter |
| *Gal* | 0.002 | -2.7 | Galanin And GMAP Prepropeptide |
| *Dgcr8* | 0.002 | -2.5 | DiGeorge Syndrome Critical Region 8, Microprocessor Complex Subunit |
| *Rsph1* | 0.002 | -2.4 | Radial Spoke Head 1 Homolog |
| *Ucn3* | 0.005 | -2.4 | Urocortin 3 |
| *Upk2* | 0.023 | -2.4 | Uroplakin 2 |
| *Oxt* | 0.002 | -2.1 | Oxytocin/Neurophysin I Prepropeptide |
| *Gstm1* | 0.002 | -1.9 | Glutathione S-Transferase Mu 1 |
| *Cox4i2* | 0.002 | -1.9 | Cytochrome C Oxidase (COX) Subunit 4 Isoform 2, Mitochondrial |
| *Dad1* | 0.002 | -1.8 | Defender Against Cell Death 1 |
| *Sept1* | 0.005 | -1.8 | Septin 1 / Differentiation 6 |
| *Gstm6* | 0.002 | -1.8 | Glutathione S-Transferase Mu 6 |
| *Ercc2* | 0.003 | -1.7 | Excision Repair Cross-Complementation Group 2 |
| *Tekt4* | 0.002 | -1.7 | Tektin 4 |
| *Pvalb* | 0.002 | -1.7 | Parvalbumin Alpha |
| *Ocm2* | 0.003 | -1.7 | Oncomodulin 2 / Parvalbumin Beta |
| *ApoE* | 0.002 | -1.6 | Apolipoprotein E / Alzheimer Disease 2 (APOE*E4-Associated, Late Onset) |
| *Uqcrb* | 0.002 | -1.6 | Ubiquinol-Cytochrome C Reductase Binding Protein |
| *Crip1* | 0.002 | -1.6 | Cysteine-Rich Intestinal Protein |
| *Ghrh* | 0.002 | -1.6 | Growth Hormone Releasing Hormone |
| *Rax* | 0.039 | -1.6 | Retina And Anterior Neural Fold Homeobox |

To comprehensively identify the functions associated with the 2,464 transcripts, gene ontology (GO) and pathways analyses were performed (Figure 3D). Gene ontology analysis revealed a significant (*q*<0.05) enrichment for the biological processes of translation, protein localization to organelle, and response to oxidative stress. Mitochondrial membrane, focal adhesion, and synapse part were the significantly (*q*<0.05) enriched cellular components. Structural constituent of ribosome, peptidase activator activity, and oxidoreductase activity acting on NAD(P)H were the most unique significant (*q*<0.05) molecular functions. Pathway analyses revealed a significant (*q*<0.05) enrichment for eIF2 signaling, regulation of eIF4 and p760K signaling, oxidative phosphorylation, mitochondrial dysfunction, and mTOR signaling (Figure 3D). Notably, most of the ontologies and pathways are related to the output of PI3K/AKT/mTOR/PTEN signaling^34^, particularly translation by the ribosome (eIF2 and eIF4 signaling) and oxidative phosphorylation in the mitochondria. Motif enrichment analysis identified several transcription factor binding sites, with the top significant (*q*<0.0001) enrichment representing binding sites for the transcription factor Elk1 that were present in 31% of transcripts (**Supplemental Table 7**). Elk1 is part of an insulin signaling network that cross-talks with PI3K/AKT/mTOR/PTEN signaling^34^ and is also involved in synaptic plasticity, learning, and neurodegenerative disease^35^. Together, these results detail differential regulation of a variety of cellular processes, and signaling pathways not previously associated with AMC, which are consistent with neurodevelopmental events occurring in PND9 hypothalamus during AMC^34^.

To validate and expand on DE genes identified by genome-wide RNA-seq analysis, reverse transcriptase quantitative PCR (RT-qPCR) analysis was performed on select genes of interest in both males and females as well as a second, independent cohort of AMC offspring. RT-qPCR analysis confirmed that *Crh* (**Supplemental Figure 5A**) was significantly reduced in AMC hypothalamus (*p*=0.0006). There was a trend (*p*=0.07) for an increased expression of *Ube3a* ligase domain containing isoforms (**Supplemental Figure 5B**), which were discovered using RNA-seq. There was also a significant (*p*<0.0001) reduction in transcripts containing the alternative 3’UTR of *Ube3a1* (**Supplemental Figure 5C**), which has the non-coding function of a competing endogenous RNA (ceRNA) or “miRNA sponge” that is critical to neurodevelopment^36^.

### Reciprocal differences in miRNA and target gene expression

One of the top gene expression changes was decreased expression of the *Dgcr8* microprocessor complex subunit gene (Table 2). Dgcr8, in combination with Drosha, is involved in the processing of primary miRNA (pri-miRNA) transcripts into precursor miRNAs (pre-miRNA)^37^. The decreased expression of *Dgcr8* suggests alterations to miRNA processing. Also suggestive of alterations to miRNA regulation was the decrease in the *Ube3a1* ceRNA. Since the RNA-seq data from PND9 hypothalamus did not accurately profile miRNA transcripts, we next evaluated the impact of AMC on miRNA expression using a multiplex miRNA expression assay on PND9 hypothalamus from handled and control pups. A total of 156 miRNAs were expressed at levels above background, with five showing significant (*p*<0.05) differential expression between AMC and control samples. Of these five, transcript levels for *rno-miR-488*, *rno-miR-144,* and *rno-miR-542-5p* were increased while transcript levels for *rno-miR-421* and *rno-miR-376b-5p* were reduced (Figure 4A). The five significant (*p*<0.05) DE miRNAs were then analyzed alongside the significant (*q*<0.05) DE mRNAs for both a bioinformatically predicted match and reciprocal DE relationship in order to produce a list of 127 putative target genes (**Supplemental Table 8**). Among the 127 putative mRNA targets of DE miRNAs are the stress response genes *Gal (rno-miR-144)*, *Pomc (rno-miR-488),* and *Pvalb (rno-miR-488)*^38–40^. Other miRNA targets include the sialidase *Neu1 (rno-miR-542-5p)* and the Amyloid beta precursor protein (APP) protease *Adam10 (rno-miR-421)*, and transcripts for the neuronal regulators *Pcdh9 (rno-miR-376b-5p)* and *Avpr1a (rno-miR-488*) (**Supplemental Table 8**)^41–43^. Finally, *rno-mir-542-5p* was predicted to target the *Ube3a1* ceRNA and the two showed a reciprocal relationship in expression (Figure 4B).

**Figure 4.**
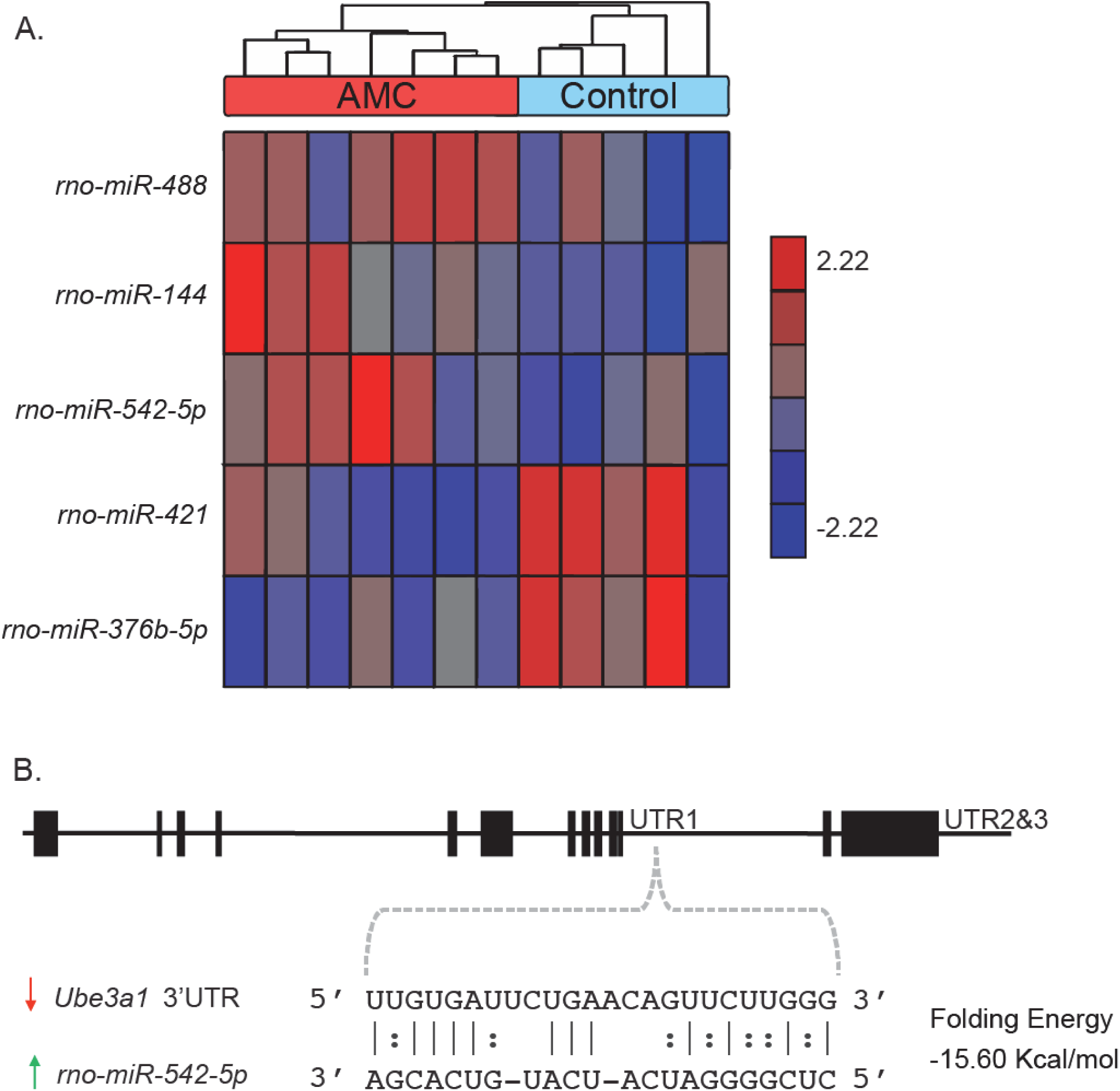
Differences in miRNA expression between AMC (*n*=7) and control (*n*=5) PND9 hypothalami. (A) Hierarchal clustering analysis of significant (*p*<0.05) differential miRNA expression between. The expression of each gene was normalized to a mean of 0 and a standard deviation of 1. The values range from −2.22 to 2.22. Down-regulated genes with AMC have negative values (blue), genes with no change have a value of zero (grey), and up-regulated genes with AMC have a positive value (red). (B) Base-pairing of the predicted relationship between *rno-mir-542-5p* and the *Ube3a1* 3’UTR.

### Integrative epigenomic prioritization of differentially regulated genes

To identify a convergent set of genes regulated by AMC in hypothalamus we integrated genes identified by the DNA methylation, RNA-seq, and multiplex miRNA assay analyses. For this combined analysis the 127 unique genes potentially regulated by the five differential miRNAs from **Supplemental Table 8** were compared with the 4,023 unique genes in proximity to DMRs shown in Figure 2. The overlap between these three gene lists resulted in 20 unique genes (Figure 5A). The presence of factors from diverse processes suggests that many pathways contribute to the AMC response in hypothalamus (Table 3). For example, *Avrp1a* functions in the stress pathway and regulates social behaviors, *Kcnh5* is an ion channel that regulates hormone and neurotransmitter release, and *Ring1* is representative of several factors on this list that repress gene expression levels (Table 3)^44–46^. *Ring1*, a component of the Polycomb Repressive Complex 1 (PRC1) complex, together with *Ube3a*, another ubiquitin E3 ligase, has been recently shown to regulate genes associated with autism^23^. The integration of genome-wide RNA-seq, DMRs, and miRNA target data sets produces a refined set of gene transcripts (Figure 5A), while also demonstrating the genome-wide nature of the differential expression and epigenetic modification that follows AMC (Figure 5B).

**Figure 5.**
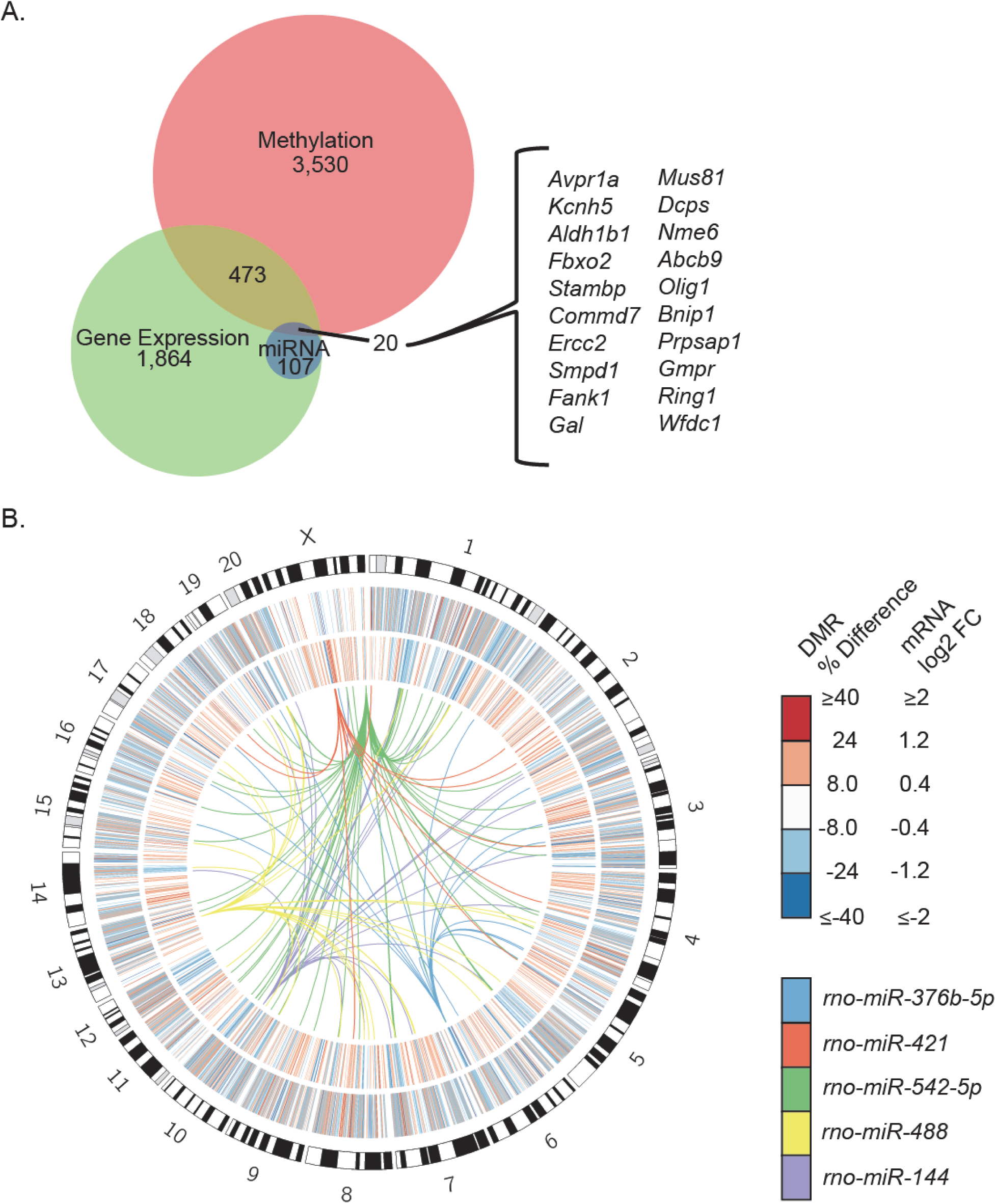
AMC is associated with genome-wide differences in DNA methylation and gene expression. (A) Integration of DMR, RNA-seq, and miRNA analyses yields 20 prioritized genes (inset). (B) Circos plot of the different data sets. The outer heatmap represents the percent differences in methylation of DMRs, the inner heatmap represents the log2 fold change of DE transcripts, and the links represent miRNAs and their predicted target genes.

**Table 3.**
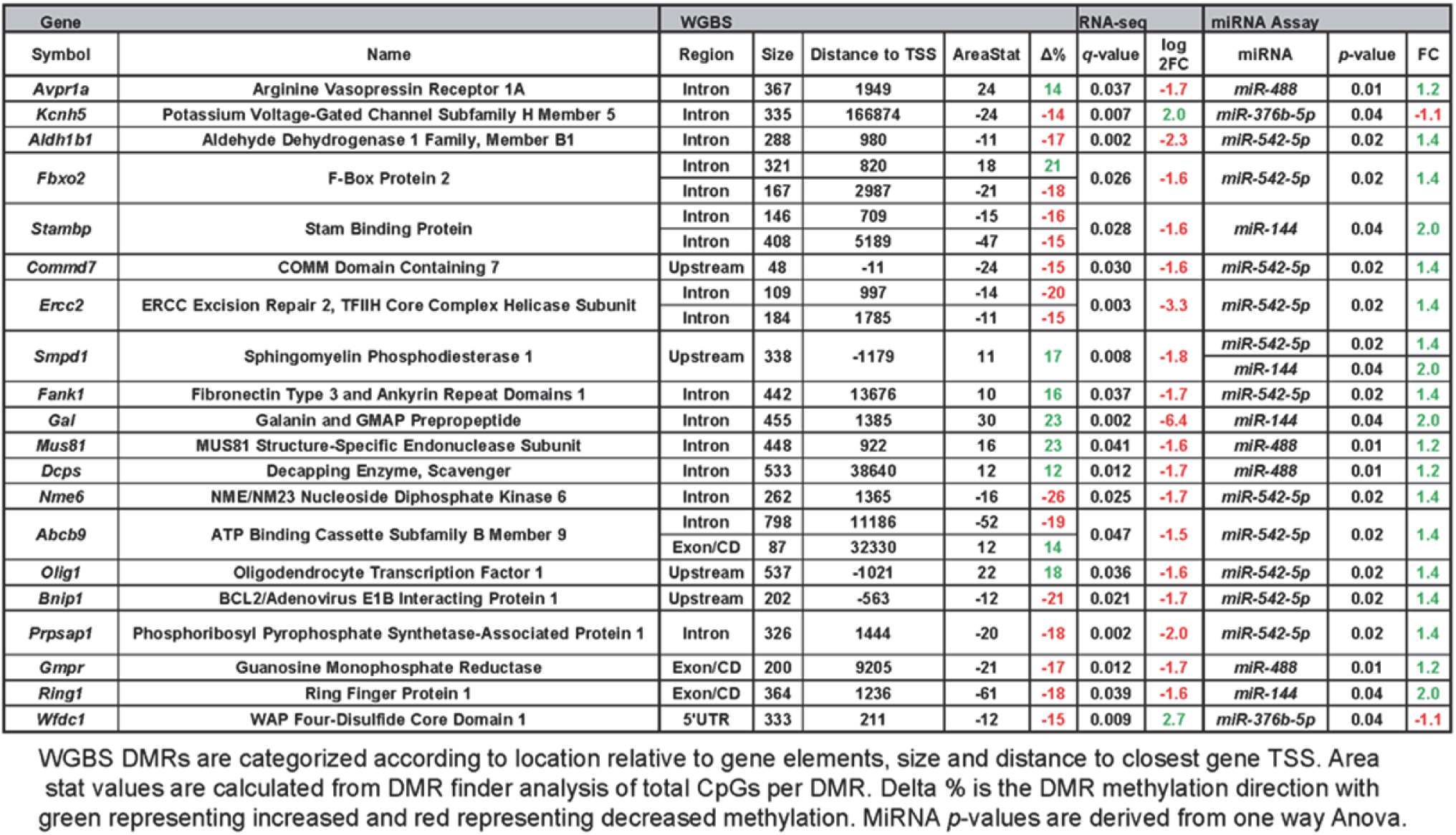
Intefrated DMR and miRNA analysis.

## Discussion

Early life environment has tremendous power to shape neurodevelopment with long-term consequences^1–3^. In humans, disjointed or reduced maternal care that occurs in cases of poverty, maternal depression, or addiction is associated with increased risk for developing cognitive and emotional deficits, neuropsychiatric disorders, and cognitive decline later in life^47,48^. On the opposite end of the spectrum, studies in humans suggest that a positive early-life environment can produce resilience to affective disorders^49–51^. In animal models, enhancements in maternal care can positively impact the developing brain leading to reduced anxiety, resilience to depression-like behaviors, and enhanced learning and memory^10,11,15^. The re-programming of the stress response in offspring receiving augmented maternal care can last a lifetime^52^ and help offset age-related cognitive deficits^14^. Much of the previous work on AMC has focused on regulation of stress response genes such as *Crh* and the *Nr3c1*. While critical for the AMC phenotype^11^ these two components of the stress system are likely not the only important factors for such large-scale and diverse reprogramming of the developing brain.

Together, our results provide the first integrative, genome-wide overview of the genes and pathways that contribute to the “reprogrammed” cognitive and emotional phenotype resulting from AMC. The results offer five novel observations. First, we identified a significant (*p*=0.015) brain region specific reduction of global DNA methylation in the hypothalamus as well as five large-scale PMDs. Second, we identified 9,439 significant (*p*<0.05) small-scale DMRs associated with 4,023 unique genes, which function in neurodevelopment and neurotransmission and may contribute to the establishment and maintenance of the behavioral phenotype. Third, we identified 2,464 significant (*q*<0.05) DE transcripts by RNA-seq analysis that extends our understanding of the gene expression differences to neurotransmission and other crucial neurologic functions by highlighting a role for translation and oxidative phosphorylation. Fourth, we identified the predicted targets of 5 significantly (*p*<0.05) altered miRNAs and also a putative interaction between *mir-542-5p* and the *Ube3a1* 3’UTR. Finally, we integrated these diverse data sets and identified a list of 20 genes with three lines of evidence (methylation, expression, and altered miRNA target) that are highly relevant to the stress resiliency phenotype resulting from AMC.

Consistent with prior descriptions for reduced methylation together with reduced CRH expression in the hypothalamus of AMC rats^21^, the connection between DNA methylation and gene expression in our genome-wide study was complex, and more apparent at the level of pathways than individual genes. At the level of gene ontology, the cellular component term “synapse part” is significantly (*q*<0.05) enriched in both datasets. Reactive oxygen species (ROS) generated by oxidative phosphorylation in the mitochondria are not only a key signaling molecule behind synaptic plasticity and memory formation^53^, but are also critical to neurodevelopment since they regulate PI3K/AKT/mTOR/PTEN signaling^54^. Furthermore, the hypomethylated *Grb10* gold DMR (FWER < 0.05) belongs to a genomically-imprinted gene encoding for a receptor-binding protein that interacts with insulin and insulin-like growth factor receptors of the PI3K/AKT/mTOR/PTEN signaling pathway^55^. Translational control is achieved via eukaryotic initiation factors (eIFs) that interact with the ribosome to drive the protein synthesis needed for synaptic plasticity^56^. Additionally, PI3K/AKT/mTOR/PTEN signaling, Wnt/β-catenin signaling, and protein kinase A (PKA) signaling converge to shape neuroplasticity, neurogenesis, cell survival, and resilience^57^. Finally, protein kinase A signaling is activated by G-protein-coupled receptors (GPCRs) that have neurotransmitter and hormone ligands, which include CRH and URN3^58^. The GPCRs of PKA signaling utilize cyclic adenosine monophosphate (cAMP) as a secondary messenger to activate PKA, which is also affected by handling to alter relevant gene expression profiles, potentially including the glucocorticoid receptor gene *Nr3c1*^52^. Ultimately, the overlapping pathways between the methylation and gene expression suggest a broad and potentially long-term impact on numerous cellular and molecular functions important for hypothalamic function.

The genes and pathways identified from our unbiased analysis of AMC-induced hypothalamic changes may be used towards the development of biomarkers and therapeutics. Suderman *et al.* have shown that epigenetic programming of the HPA axis by maternal care is conserved in humans at a select locus^59^. Given the potential conservation, the suite of new candidate loci identified may be examined in humans towards the development of novel epigenetic biomarkers of resiliency^60^. Such a diagnostic may then be used to identify individuals that would benefit from corrective measures, which include environmental enrichment^61^ and pharmacologic agents. The suite of genetic loci identified may also be used towards the development of inventive pharmacologic agents. Other studies have shown that the transcriptomic profile and epigenetic modifications of altered maternal care can be reprogrammed at adulthood by the global epigenetic modifiers, specifically the methyl donor L-methionine or the histone deacetylase inhibitor trichostatin A (TSA)^18,19,62,63^. More recently, it has been shown that pharmacologic agents can mimic some of the effects of AMC *in vitro*^12^. Therefore, some of the epigenomic profile established by maternal care is plastic and can be manipulated later in life. However, these previously examined modifiers result in attenuations that are global in nature and may have off-target effects. Our results could be used to overcome this limitation since the suite of therapeutic candidates identified can be utilized as prioritized targets for somatic epigenetic editing studies^64–67^. Ultimately, our results could form the molecular foundation for developing effective interventions for deficient early maternal care in humans or be developed as new treatments for psychiatric disorders^67^.

## Acknowledgements

This work used the Vincent J. Coates Genomics Sequencing Laboratory at UC Berkeley, supported by NIH [S10 OD018174] Instrumentation Grant. A.V.C. was supported by National Institutes of Health [T32MH073124-06]. B.I.L. was supported by a postdoctoral fellowship award from the Canadian Institutes of Health Research (CIHR). A.S-T and T.Z.B. were supported by NIH MH73136 and P50 096889.

## References

1 Levine S. Infantile experience and resistance to physiological stress. Science 1957; 126: 405.

2 Francis D, Diorio J, Liu D, Meaney MJ. Nongenomic transmission across generations of maternal behavior and stress responses in the rat. Science 1999; 286: 1155–1158.

3 Barnett SA, Burn J. Early stimulation and maternal behaviour. Nature 1967; 213: 150–152.

4 Meaney M, Aitken D, van Berkel C, Bhatnagar S, Sapolsky R. Effect of neonatal handling on age-related impairments associated with the hippocampus. Science 1988; 239: 766–768.

5 Avishai-Eliner S, Eghbal-Ahmadi M, Tabachnik E, Brunson KL, Baram TZ. Down-regulation of hypothalamic corticotropin-releasing hormone messenger ribonucleic acid (mRNA) precedes early-life experience-induced changes in hippocampal glucocorticoid receptor mRNA. Endocrinology 2001; 142: 89–97.

6 Fenoglio KA, Chen Y, Baram TZ. Neuroplasticity of the hypothalamic-pituitary-adrenal axis early in life requires recurrent recruitment of stress-regulating brain regions. J Neurosci 2006; 26: 2434–2442.

7 Meaney MJ, Viau V, Bhatnagar S, Betito K, Iny LJ, O'Donnell D et al. Cellular mechanisms underlying the development and expression of individual differences in the hypothalamic-pituitary-adrenal stress response. J Steroid Biochem Mol Biol 1991; 39: 265–274.

8 Caldji C, Tannenbaum B, Sharma S, Francis D, Plotsky PM, Meaney MJ. Maternal care during infancy regulates the development of neural systems mediating the expression of fearfulness in the rat. Proc Natl Acad Sci U S A 1998; 95: 5335–5340.

9 Liu D, Diorio J, Tannenbaum B, Caldji C, Francis D, Freedman a et al. Maternal care, hippocampal glucocorticoid receptors, and hypothalamic-pituitary-adrenal responses to stress. Science 1997; 277: 1659–1662.

10 Viau V, Sharma S, Plotsky P, Meaney M. Increased plasma ACTH responses to stress in nonhandled compared with handled rats require basal levels of corticosterone and are associated with increased levels of ACTH secretagogues in the median eminence. J Neurosci 1993; 13: 1097–1105.

11 Plotsky PM, Meaney MJ. Early, postnatal experience alters hypothalamic corticotropin-releasing factor (CRF) mRNA, median eminence CRF content and stress-induced release in adult rats. Mol Brain Res 1993; 18: 195–200.

12 Singh-Taylor A, Molet J, Jiang S, Korosi A, Bolton JL, Noam Y et al. NRSF-dependent epigenetic mechanisms contribute to programming of stress-sensitive neurons by neonatal experience, promoting resilience. Mol Psychiatry 2017. doi:10.1038/mp.2016.240.

13 Fenoglio KA, Brunson KL, Avishai-Eliner S, Stone BA, Kapadia BJ, Baram TZ. Enduring, Handling-Evoked Enhancement of Hippocampal Memory Function and Glucocorticoid Receptor Expression Involves Activation of the Corticotropin-Releasing Factor Type 1 Receptor. Endocrinology 2005; 146: 4090–4096.

14 Korosi A, Baram TZ. The pathways from mother's love to baby's future. Front Behav Neurosci 2009; 3: 27.

15 Meaney MJ, Aitken DH. The effects of early postnatal handling on hippocampal glucocorticoid receptor concentrations: temporal parameters. Dev Brain Res 1985; 22: 301–304.

16 Korosi A, Shanabrough M, McClelland S, Liu Z-W, Borok E, Gao X-B et al. Early-Life Experience Reduces Excitation to Stress-Responsive Hypothalamic Neurons and Reprograms the Expression of Corticotropin-Releasing Hormone. J Neurosci 2010; 30: 703–713.

17 Singh-Taylor A, Korosi A, Molet J, Gunn BG, Baram TZ. Synaptic rewiring of stress-sensitive neurons by early-life experience: A mechanism for resilience? Neurobiol. Stress. 2015; 1: 109–115.

18 Weaver ICG, Cervoni N, Champagne FA, D'Alessio AC, Sharma S, Seckl JR et al. Epigenetic programming by maternal behavior. Nat Neurosci 2004; 7: 847–854.

19 Weaver ICG, Meaney MJ, Szyf M. Maternal care effects on the hippocampal transcriptome and anxiety-mediated behaviors in the offspring that are reversible in adulthood. Proc Natl Acad Sci U S A 2006; 103: 3480–3485.

20 McGowan PO, Suderman M, Sasaki A, Huang TCT, Hallett M, Meaney MJ et al. Broad Epigenetic Signature of Maternal Care in the Brain of Adult Rats. PLoS One 2011; 6: e14739.

21 McClelland S, Korosi A, Cope J, Ivy A, Baram TZ. Emerging roles of epigenetic mechanisms in the enduring effects of early-life stress and experience on learning and memory. Neurobiol Learn Mem 2011; 96: 79–88.

22 Schroeder DI, Lott P, Korf I, LaSalle JM. Large-scale methylation domains mark a functional subset of neuronally expressed genes. Genome Res 2011; 21: 1583–1591.

23 Dunaway KW, Islam MS, Coulson RL, Lopez SJ, Vogel Ciernia A, Chu RG et al. Cumulative Impact of Polychlorinated Biphenyl and Large Chromosomal Duplications on DNA Methylation, Chromatin, and Expression of Autism Candidate Genes. Cell Rep 2016; 17: 3035–3048.

24 Lister R, Mukamel EA, Nery JR, Urich M, Puddifoot CA, Johnson ND et al. Global Epigenomic Reconfiguration During Mammalian Brain Development. Science 2013; 341: 1237905.

25 Geraldo S, Khanzada UK, Parsons M, Chilton JK, Gordon-Weeks PR. Targeting of the F-actin-binding protein drebrin by the microtubule plus-tip protein EB3 is required for neuritogenesis. Nat Cell Biol 2008; 10: 1181–1189.

26 Murata Y, Doi T, Taniguchi H, Fujiyoshi Y. Proteomic analysis revealed a novel synaptic proline-rich membrane protein (PRR7) associated with PSD-95 and NMDA receptor. Biochem Biophys Res Commun 2005; 327: 183–191.

27 Brunet A, Bonni A, Zigmond MJ, Lin MZ, Juo P, Hu LS et al. Akt Promotes Cell Survival by Phosphorylating and Inhibiting a Forkhead Transcription Factor. Cell 1999; 96: 857–868.

28 Houge G, Rasmussen IH, Hovland R. Loss-of-Function CNKSR2 Mutation Is a Likely Cause of Non-Syndromic X-Linked Intellectual Disability. Mol Syndromol 2012; 2: 60–63.

29 Lee SR, Ramos SM, Ko A, Masiello D, Swanson KD, Lu ML et al. AR and ER interaction with a p21-activated kinase (PAK6). Mol Endocrinol 2002; 16: 85–99.

30 Hansen KD, Langmead B, Irizarry RA. BSmooth: from whole genome bisulfite sequencing reads to differentially methylated regions. Genome Biol 2012; 13: R83.

31 Ong C-T, Corces VG. CTCF: an architectural protein bridging genome topology and function. Nat Publ Gr 2014; 15: 234–246.

32 Antinucci P, Nikolaou N, Meyer MP, Hindges R. Teneurin-3 specifies morphological and functional connectivity of retinal ganglion cells in the vertebrate visual system. Cell Rep 2013; 5: 582–92.

33 Ma H, Ng HM, Teh X, Li H, Lee YH, Chong YM et al. Zfp322a Regulates Mouse ES Cell Pluripotency and Enhances Reprogramming Efficiency. PLoS Genet 2014; 10: e1004038.

34 Nyman E, Rajan MR, Fagerholm S, Brännmark C, Cedersund G, Strålfors P. A Single Mechanism Can Explain Network-wide Insulin Resistance in Adipocytes from Obese Patients with Type 2 Diabetes. 2014; 289: 33215–33230,.

35 Besnard A, Galan-Rodriguez B, Vanhoutte P, Caboche J. Elk-1 a Transcription Factor with Multiple Facets in the Brain. Front Neurosci 2011; 5: 35.

36 Valluy J, Bicker S, Aksoy-Aksel A, Lackinger M, Sumer S, Fiore R et al. A coding-independent function of an alternative Ube3a transcript during neuronal development. Nat Neurosci 2015; 18: 666–673.

37 Han J, Lee Y, Yeom K-H, Nam J-W, Heo I, Rhee J-K et al. Molecular Basis for the Recognition of Primary microRNAs by the Drosha-DGCR8 Complex. Cell 2006; 125: 887–901.

38 Pritchard LE, Turnbull A V, White A. REVIEW Pro-opiomelanocortin processing in the hypothalamus: impact on melanocortin signalling and obesity. J Endocrinol 2002; 172: 411–421.

39 Khoshbouei H, Cecchi M, Dove S, Javors M, Morilak DA. Behavioral reactivity to stress: amplification of stress-induced noradrenergic activation elicits a galanin-mediated anxiolytic effect in central amygdala. Pharmacol Biochem Behav 2002; 71: 407–417.

40 Hu W, Zhang M, Czéh B, Flügge G, Zhang W. Stress Impairs GABAergic Network Function in the Hippocampus by Activating Nongenomic Glucocorticoid Receptors and Affecting the Integrity of the Parvalbumin-Expressing Neuronal Network. Neuropsychopharmacology 2010; 35: 1693–1707.

41 Lammich S, Kojro E, Postina R, Gilbert S, Pfeiffer R, Jasionowski M et al. Constitutive and regulated alpha-secretase cleavage of Alzheimer's amyloid precursor protein by a disintegrin metalloprotease. Proc Natl Acad Sci U S A 1999; 96: 3922–3927.

42 Marshall CR, Noor A, Vincent JB, Lionel AC, Feuk L, Skaug J et al. Structural variation of chromosomes in autism spectrum disorder. Am J Hum Genet 2008; 82: 477–488.

43 Murgatroyd C, Spengler D. Epigenetic programming of the HPA axis: early life decides. Stress 2011; 14: 581–589.

44 Pappa I, St Pourcain B, Benke K, Cavadino A, Hakulinen C, Nivard MG et al. A genome-wide approach to children's aggressive behavior: The EAGLE consortium. Am J Med Genet Part B Neuropsychiatr Genet 2016; 171: 562–572.

45 Ju M, Wray D. Molecular identification and characterisation of the human eag2 potassium channel. FEBS Lett 2002; 524: 204–210.

46 Satijn DP, Gunster MJ, van der Vlag J, Hamer KM, Schul W, Alkema MJ et al. RING1 is associated with the polycomb group protein complex and acts as a transcriptional repressor. Mol Cell Biol 1997; 17: 4105–4113.

47 Lupien SJ, King S, Meaney MJ, McEwen BS. Child's stress hormone levels correlate with mother's socioeconomic status and depressive state. Biol Psychiatry 2000; 48: 976–980.

48 Baram TZ, Davis EP, Obenaus A, Sandman CA, Small SL, Solodkin A et al. Fragmentation and unpredictability of early-life experience in mental disorders. Am. J. Psychiatry. 2012; 169: 907–915.

49 Gunnar MR, Frenn K, Wewerka SS, Van Ryzin MJ. Moderate versus severe early life stress: associations with stress reactivity and regulation in 10-12-year-old children. Psychoneuroendocrinology 2009; 34: 62–75.

50 Halligan SL, Herbert J, Goodyer I, Murray L. Disturbances in morning cortisol secretion in association with maternal postnatal depression predict subsequent depressive symptomatology in adolescents. Biol Psychiatry 2007; 62: 40–46.

51 Wilson RS, Schneider JA, Boyle PA, Arnold SE, Tang Y, Bennett DA. Chronic distress and incidence of mild cognitive impairment. Neurology 2007; 68: 2085–2092.

52 Gourion D, Arseneault L, Vitaro F, Brezo J, Turecki G, Tremblay RE. Early environment and major depression in young adults: a longitudinal study. Psychiatry Res 2008; 161: 170–176.

53 Kishida KT, Klann E. Sources and Targets of Reactive Oxygen Species in Synaptic Plasticity and Memory. Antioxid Redox Signal 2006; 0: 61121054212009.

54 Ostrakhovitch EA, Semenikhin OA. The role of redox environment in neurogenic development. Arch Biochem Biophys 2013; 534: 44–54.

55 Yu Y, Yoon S-O, Poulogiannis G, Yang Q, Ma XM, Villén J et al. Phosphoproteomic Analysis Identifies Grb10 as an mTORC1 Substrate That Negatively Regulates Insulin Signaling. Science 2011; 332: 1322–1326.

56 Santini E, Huynh TN, Klann E. Mechanisms of Translation Control Underlying Long-Lasting Synaptic Plasticity and the Consolidation of Long-Term Memory. In: Progress in molecular biology and translational science. 2014, pp 131–167.

57 Charney DS, Manji HK. Life Stress, Genes, and Depression: Multiple Pathways Lead to Increased Risk and New Opportunities for Intervention. Sci Signal 2004; 2004: re5.

58 A G Henckens MJ, Deussing JM, Chen A. Region-specific roles of the corticotropin-releasing factor-urocortin system in stress. 2016; 17: 636–651.

59 Suderman M, McGowan PO, Sasaki A, Huang TCT, Hallett MT, Meaney MJ et al. Conserved epigenetic sensitivity to early life experience in the rat and human hippocampus. Proc Natl Acad Sci U S A 2012;: 17266–17272.

60 consortium TB. Quantitative comparison of DNA methylation assays for biomarker development and clinical applications. Nat Biotech 2016; 34: 726–737.

61 Francis DD, Diorio J, Plotsky PM, Meaney MJ. Environmental enrichment reverses the effects of maternal separation on stress reactivity. J Neurosci 2002; 22: 7840–7843.

62 Champagne FA, Francis DD, Mar A, Meaney MJ. Variations in maternal care in the rat as a mediating influence for the effects of environment on development. Physiol Behav 2003; 79: 359–371.

63 Weaver ICG, Champagne FA, Brown SE, Dymov S, Sharma S, Meaney MJ et al. Reversal of Maternal Programming of Stress Responses in Adult Offspring through Methyl Supplementation: Altering Epigenetic Marking Later in Life. J Neurosci 2005; 25: 11045–11054.

64 Laufer BI, Singh SM. Strategies for precision modulation of gene expression by epigenome editing: an overview. Epigenetics Chromatin 2015; 8: 34.

65 Stelzer Y, Wu H, Song Y, Shivalila CS, Markoulaki S, Jaenisch R. Parent-of-Origin DNA Methylation Dynamics during Mouse Development. Cell Rep 2016; 16: 3167–3180.

66 Morita S, Noguchi H, Horii T, Nakabayashi K, Kimura M, Okamura K et al. Targeted DNA demethylation in vivo using dCas9-peptide repeat and scFv-TET1 catalytic domain fusions. Nat Biotechnol 2016; 34: 1060–1065.

67 Stepper P, Kungulovski G, Jurkowska RZ, Chandra T, Krueger F, Reinhardt R et al. Efficient targeted DNA methylation with chimeric dCas9-Dnmt3a-Dnmt3L methyltransferase. Nucleic Acids Res 2017; 45: 1703–1713.

